# Antagonizing the SAMD9 pathway is key to myxoma virus host shut-off and immune evasion

**DOI:** 10.1101/2024.02.01.578447

**Authors:** Tahseen Raza, Erich Perterson, Jason Liem, Jia Liu

## Abstract

Poxviruses have life cycles exclusively in the cytoplasm. However, these viruses can have profound impact to host transcription. One possible mechanism is through viral manipulation of host protein synthesis and such ability is critical for viral immune evasion. Many mammalian poxviruses encode more than one viral protein to interact with the host SAMD9 protein. In myxoma virus (MYXV), a rabbit specific poxvirus and non-pathogenic for other species, viral M062 protein is the lone inhibitor to SAMD9 with broad species specificity and loss of *M062R* in viral genome (Δ*M062R* mutant) leads to profound infection defect. We previously found Δ*M062R* remodeled transcriptomic landscape in monocytes/macrophages that is associated with the crosstalk between the SAMD9 pathway and cGAS/STING/IRF3 DNA sensing pathway. In this study we completed the characterization of Δ*M062R* infection. We observed that although this replication-defective virus preserved intact early protein synthesis, it failed to conduct host shutoff. Despite a defect in viral DNA replication, Δ*M062R* infection retained intact intermediate protein synthesis comparable to the wildtype virus. Using time course dual RNAseq analyses we found that the overall viral gene transcription profile was mostly indistinguishable from that of the wildtype MYXV. However, the slightly attenuated late RNA synthesis along with the block at viral protein synthesis led to its infection defect. Infection by Δ*M062R* in macrophages potentiated the antiviral responses to new danger signals. We provided an initial characterization of such a state in which host antiviral protein synthesis may be promoted leading to the immunological consequence.

**Importance:** Poxviruses utilize multi-faceted strategies to evade and manipulate host immunity. Through targeted gene deletion, we generated useful tools of mutant poxviruses to investigate specific crosstalk between host defense mechanisms. Through studying MYXV M062 protein function, we previously identified SAMD9 as one host target of poxvirus *C7L* superfamily, in which family *M062R* is one member. However, what kind of cellular outcome caused by Δ*M062R* infection remained unknown. The infection defect of Δ*M062R* caused the induction of host inflammation program is likely due to the activation of the host pathway governed by SAMD9. Because little is known about SAMD9 cellular function and the pathways it regulates, which are important for cellular homeostasis and immune regulation, this study on Δ*M062R* induced effect in host cells will provide new insight on how SAMD9 affects cellular protein synthesis and immunological responses.

## Introduction

Myxoma virus (MYXV) is a poxvirus infecting lagomorph (rabbits).^1^ After MYXV spilled into European rabbits (which was not a natural host to MYXV), approximately 80 years of studies on MYXV-European rabbit co-evolution across the continents^2–4^ provides a wealth of knowledge for us to prepare for ongoing monkeypox virus co-evolution in humans.^5,6^ Studying the biology of MYXV provides insight on viral tropism determination and evasion of host immunity, both of which are highly relevant to the understanding of poxviruses capable of tropism expansion. Myxoma virus (MYXV) is also a promising virotherapeutic agent for cancer treatment^7,8^ and thus a complete understanding of its ability to regulate human cellular and immunological signaling events is urgently needed.

Myxoma virus M062 is the gene product of *M062R*, a host range determinant from poxvirus *C7L* superfamily.^9^ Without *M062R*, the resulting mutant virus loses its ability of productive infection in almost all cells tested and *in vivo*.^10^ One host target of viral M062 is SAMD9,^10,11^ a large cytoplasmic protein with a complex domain structure^12^ and ability of cleaving tRNA^phe^.^13^ Deleterious mutations in SAMD9 often lead to hematological disorders (often also involving myeloid cells),^14,15^ inflammatory disorder,^16,17^ and diseases with complex syndromes during development.^18,19^ Interestingly, full-length SAMD9 can be associated with dsDNA with and without viral infection, and the presence of MYXV M062 prevents it from binding to dsDNA.^20^ The behavior of SAMD9 recognizing dsDNA possibly is in part due to the AlbA_2-like domain (134-385 amino acids of SAMD9) that can bind to double-stranded nucleic acids in a sequence non-specific manner.^21^ The replication defective MYXV with targeted *M062R* gene induces a unique inflammatory state through the activation of both cGAS/STING/IRF3 DNA sensing axis and SAMD9 pathway.^20^ In this study, we further delineated the infection defect of *M062R*-null MYXV (Δ*M062R*), characterized Δ*M062R* infection caused outcome on viral and host protein translation, and reported the effect of Δ*M062R* in potentiating macrophages in response to new danger signals. We also summarized the effect on the transcriptomic landscape in monocytes/macrophages induced by Δ*M062R* and concluded that these characteristics of Δ*M062R* may be key to its immunotherapeutic potential in reversing the immunosuppressive environment such as the tumor environment.

## Results

### *M062R*-null MYXV (Δ*M062R*) caused abortive infection and failed to conduct host shutoff

We previously found that Δ*M062R* infection caused abortive infection in almost all cells tested from species such as humans and rabbits.^10^ In human THP1 cells, a monocytic cell model system that can be differentiated into macrophage-like cells for investigation of innate immune signaling, Δ*M062R* infection also led to an abortive infection with significant reduction in viral replication compared to the wildtype MYXV (**Figure 1A**). Although early gene expression of this virus did not show significant difference from that of the wildtype MYXV,^10,20^ Δ*M062R* infection failed to lead to host shutoff (**Figure 1B**). With comparable levels of early protein synthesis during Δ*M062R* infection to WT infection, many necessary components for the next stage of viral life cycle are expected to be synthesized along with important immunoregulatory factors.

**Figure 1.**
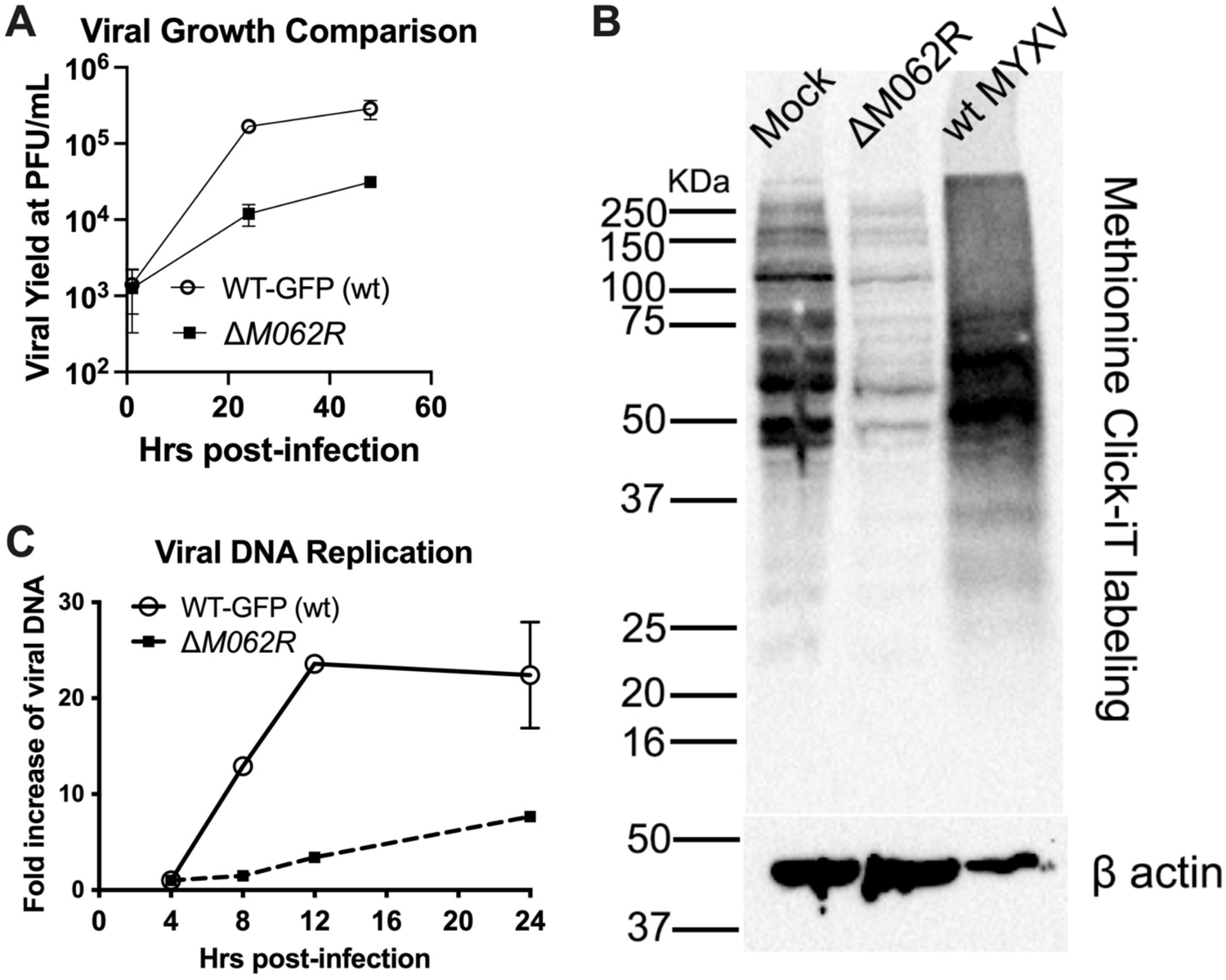
Infection by Δ*M062R* fails to conduct host shut-off with defect in viral DNA replication. A. Abortive infection by Δ*M062R* in human macrophages. THP1 cells were first differentiated into macrophages before infection with either wildtype MYXV (WT-GFP) or Δ*M062R* at an MOI of 1. At given time points (1, 24, and 48 hrs. p.i.) cell lysates were harvested for titration on BSC-40 cells. Shown is the summary of triplicate at each time point for each virus. B. Infection by Δ*M062R* cannot perform host shut-off. THP1-differentiated macrophages were mock treated, infected with Δ*M062R*, or wildtype MYXV at an MOI of 5. At 16 hrs. p.i., cells were pulse-labelled for de novo synthesized polypeptide with AHA using Click-iT chemistry. Total proteins from each group were then separated on SDS-PAGE for Western Blot. Cells from Δ*M062R* infection showed the same banding pattern as mock infected cells, while wildtype MYXV infection caused distinct pattern of protein mobility suggesting the successful shutoff of host protein synthesis and transition to predominantly viral protein synthesis. Loading of comparable quantity of total protein is indicated by β-actin levels. Shown is the representative of three independent experiments. C. DNA replication defect during Δ*M062R* infection. Cells were infected with either wildtype MYXV or Δ*M062R* at an MOI of 5. At the given time points, DNA extraction was performed followed by realtime PCR. Relative realtime PCR was performed and fold induction calculation was conducted using ΔΔCt method.

### Infection by Δ*M062R* did not affect early gene expression but significantly reduced late gene expression

Although early protein synthesis appeared intact, viral DNA replication is significantly attenuated during Δ*M062R* infection (**Figure 1C**). To evaluate overall viral gene expression, we performed RNAseq analyses in a time course to examine how viral gene expression changed with time using DESeq2Norm method.^22,23^ Despite the DNA replication defect, Δ*M062R* infection caused moderate attenuation in intermediate RNA synthesis (**Figure 2**). We next examined how intermediate protein synthesis is affected during the mutant viral infection. We utilized a well-characterized luciferase based assay with luciferase expression driven by known vaccinia virus (VACV) intermediate promoters, including that of A2L (pA2L),^24^ I1L (pI1L),^24^ and another consensus promoter sequence (pINT) previously reported by Yang et al.^25^ Shortly after infection by either wildtype or Δ*M062R* MYXV, the construct containing viral intermediate promoter driven luciferase was transfected in the absence or presence of AraC followed by measurement of luciferase expression. The presence of DNA replication inhibitor (AraC) led to reduced differences in intermediate protein expression between wildtype and Δ*M062R* infection (**Figure 3A-C**). Albeit statistically significant, moderate reduction in the intermediate protein synthesis is observed during Δ*M062R* infection (**Figure 3A-C**). The slight reduction in intermediate protein production may not be apparent using Western blot (**Figure 3D** the intermediate protein M038). Consistent with our previous finding, during Δ*M062R* infection late viral protein production is inhibited (**Figure 3D** probing of the late protein SERP1), which is preceded with reduction in viral late RNA synthesis (**Figure 4**). Although we observed apparent reduction in viral late transcripts during Δ*M062R* infection compared to wildtype infection (2-way ANOVA statistical analyses), the level differences between two infections are subtle (multiple t test comparison) (**Figure 4**). This is a highly unusual phenomenon in that the mutant virus is replication defective while capable of generating late transcripts, albeit at moderately lower levels than those from wildtype viral infection.

**Figure 2.**
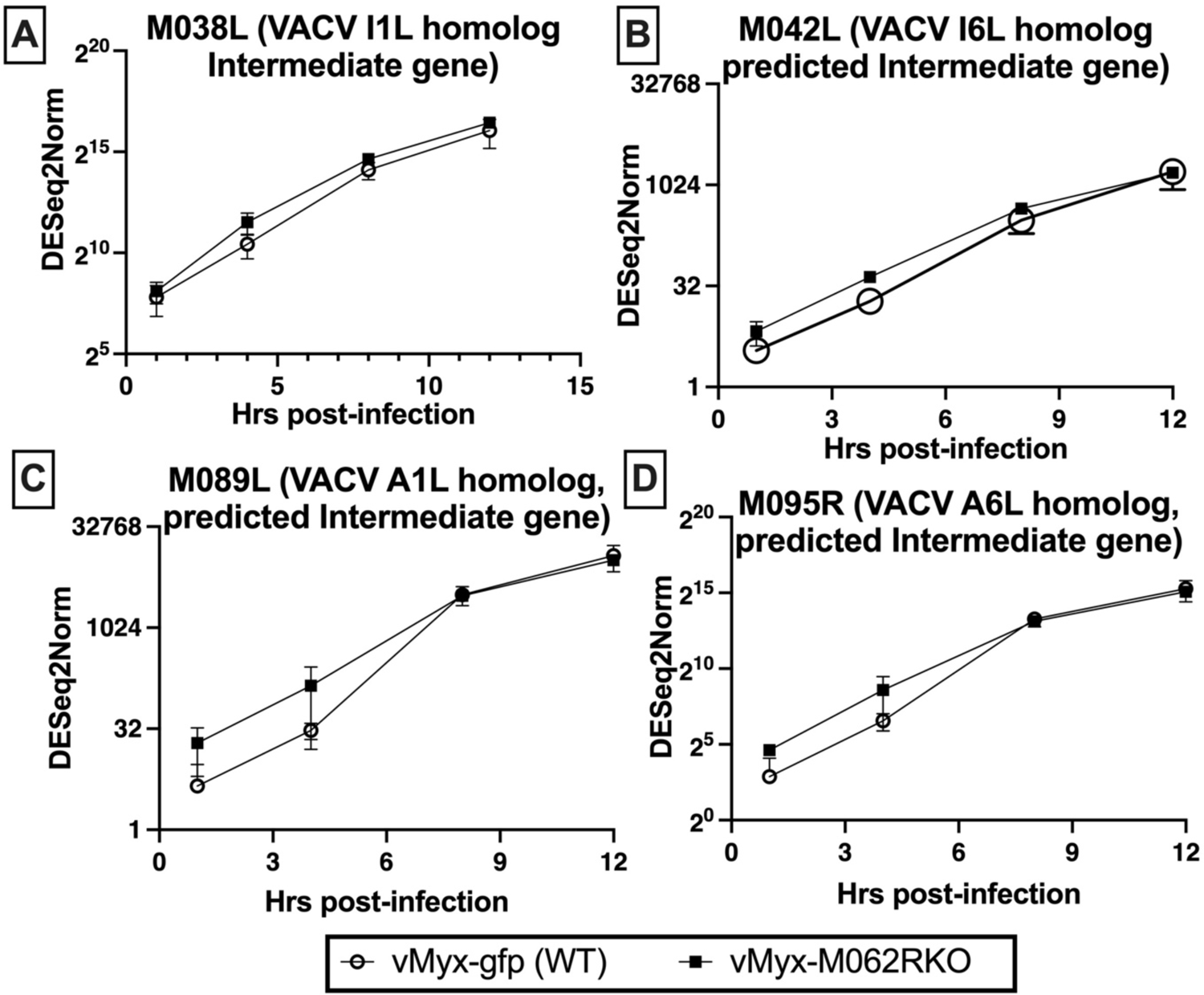
Comparison of MYXV intermediate gene transcription. RNAseq was performed using cells that were either infected with wildtype or Δ*M062R* at an MOI of 5 for a timecourse at given timepoints (1, 4, 8, and 12 hrs. p.i.). Three independent replicates were performed and shown is the summary of triplicate for each time point and each virus. Known MYXV intermediate genes (A. M038L, B. M042L, C. M089L, and D. M095R) were examined for their transcription levels throughout the time course using normalized detection quantification (the median of ratios method in DESeQ2).

**Figure 3.**
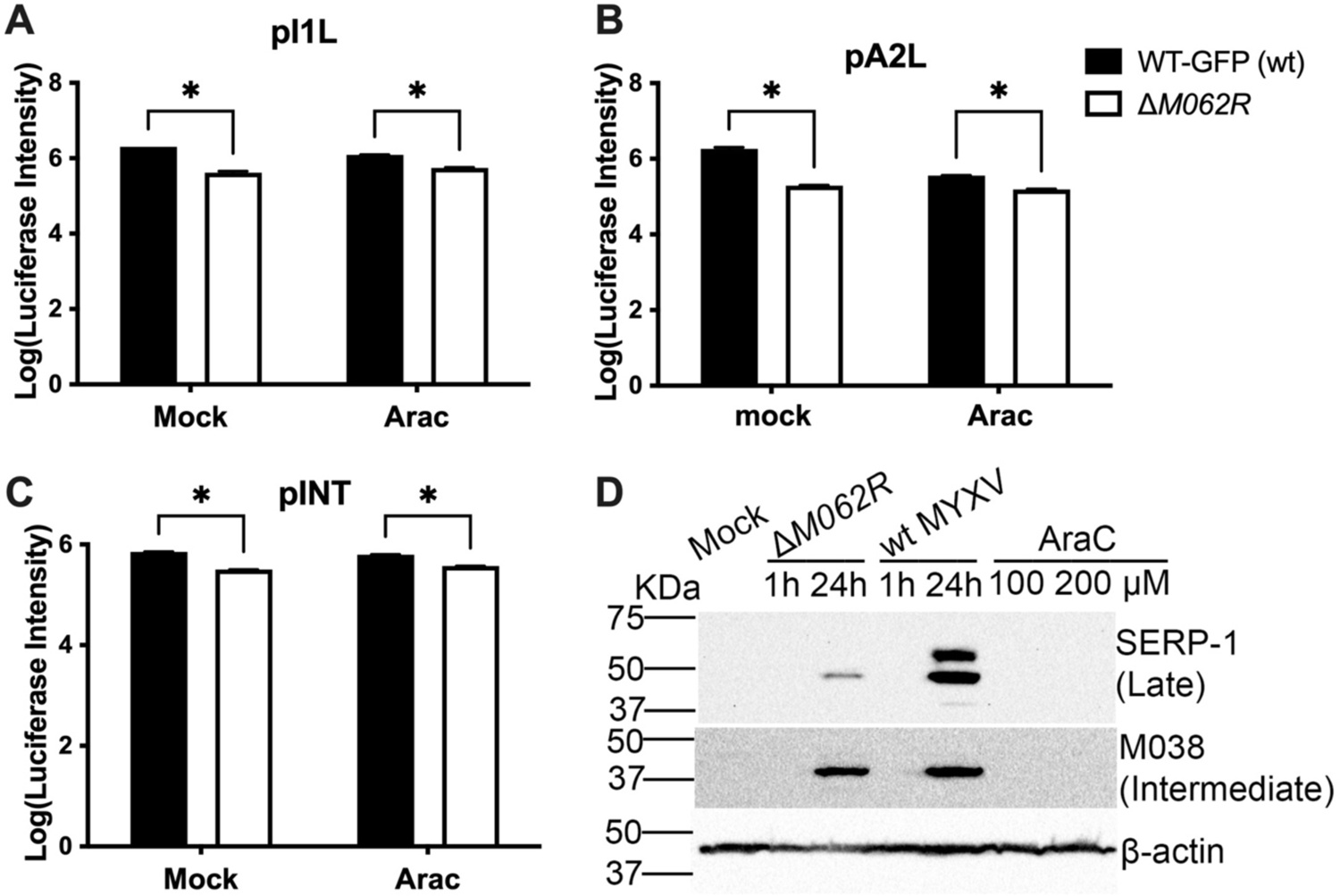
Infection by Δ*M062R* presented competent production of intermediate protein. After infection of cells with either wildtype MYXV or Δ*M062R* at an MOI of 5, in the presence or absence of DNA replication inhibitor, AraC (200 μM), transfection of viral intermediate promoter driven luciferase plasmid was performed using ViaFect, approximately 18 hrs. post transfection, supernatant were collected for luciferase acitivity assay. Results for pI1L (A), pA2L (B), and pINT (C) -driven luciferase activities are shown. Consistent results were observed using HeLa and HEK293T cells. Shown is a representative result using HeLa cells. (D) THP1 were differentiated into macrophages before mock treated, infected with Δ*M062R* or wildtype MYXV. At the given time points of 1 and 24 hrs. p.i., viral post-replicative gene expression was examined. The expression of either M038 (intermediate gene) or SERP-1 (late gene) was assessed in Western Blot. In the presence of AraC, post-replicative protein synthesis was effectively inhibited, which confirmed the kinetic group of M038 and SERP-1.

**Figure 4.**
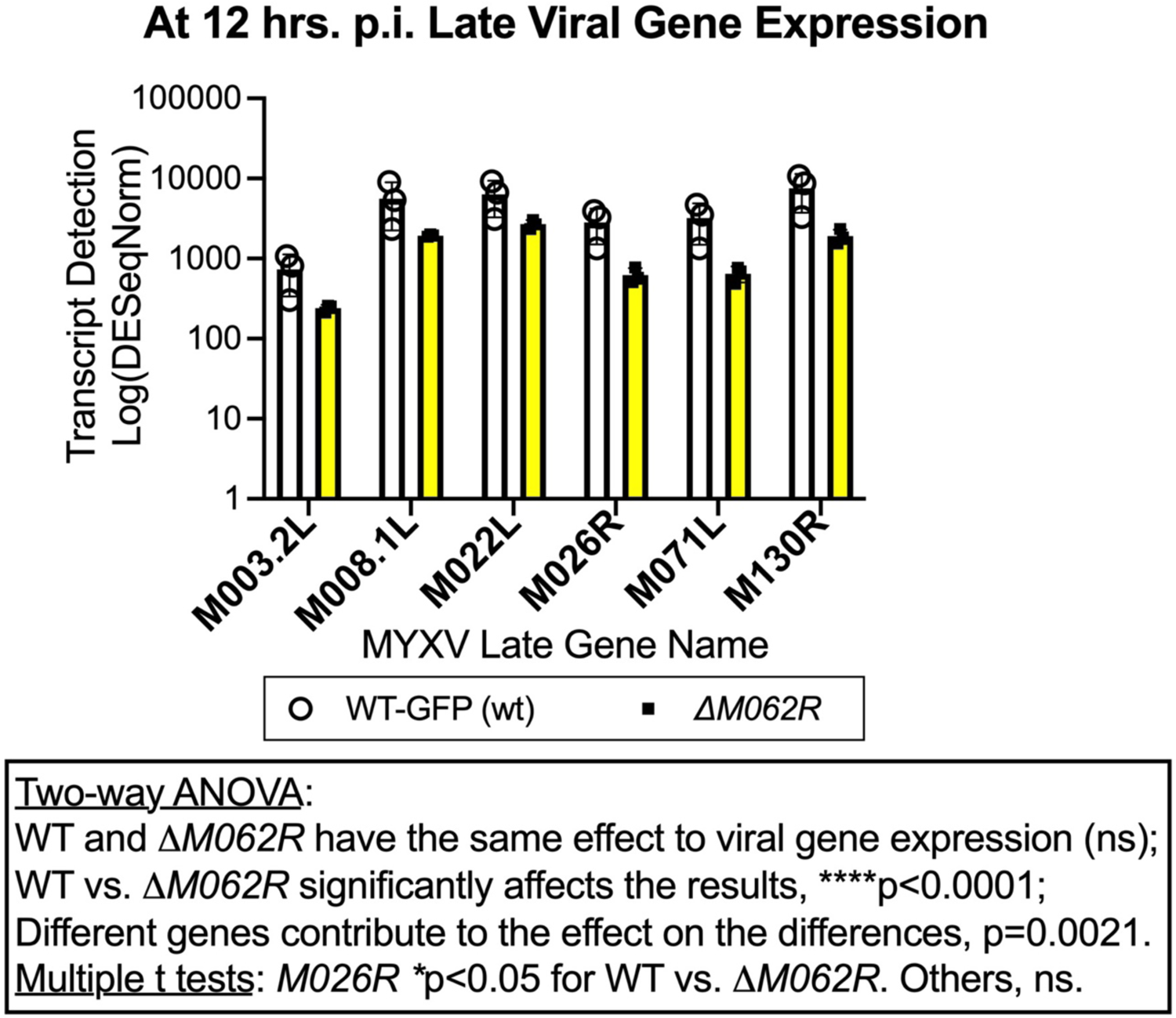
Infection by Δ*M062R* permits late gene transcription. After RNAseq data analyses, comparison of gene transcripts of known MYXV late gene transcripts at 12 hrs. p.i. between wildtype MYXV and Δ*M062R* were conducted using normalized detection quantification (the median of ratios method in DESeQ2). Triplicate of samples per viral infection were included in the study. Two-way ANOVA statistical analyses and multiple unpaired t test were performed to assess the gene expression comparison.

SAMD9 is the sole host factor in human cells causing the abortive infection by Δ*M062R* infection, as deleting SAMD9 alone restored Δ*M062R* infection to the wildtype virus levels.^11,26^ We thus concluded that during Δ*M062R* infection, un-obstructed SAMD9 might inhibit viral DNA replication (**Figure 1C**) and deplete tRNA^phe^,^13^ which gradually slowed down viral intermediate protein synthesis. This eventually blocked viral late protein synthesis. These caused failure of the mutant virus to conduct host shutoff and produce progeny virion.

### Infection by Δ*M062R* did not affect host protein synthesis, as it promoted accumulationof antiviral proteins

There are two viral proteins in VACV interacting with SAMD9, C7 and K1.^27,28^ The *C7L-K1L* double knockout VACV (VACV-Δ*C7L*Δ*K1L*) presented similar infection defect as Δ*M062R* MYXV, likely also due to SAMD9 function.^11^ Infection by VACV-Δ*C7L*Δ*K1L* induced a state of general or global translation inhibition for 5’ Cap-mRNA, IRES-containing mRNA, and viral mRNA.^29^ We initially hypothesized that infection by Δ*M062R* MYXV would trigger a similar state of global translation inhibition. However, our earlier finding of Δ*M062R* induced proinflammatory response in macrophages prompted us to rethink this hypothesis. This is because Δ*M062R* infection in human macrophages activated cellular RNA transcription and induced type 1 interferon (IFN-I) and ISGs.^20^ We thus decided to characterize the cellular translation status induced by Δ*M062R* MYXV to consolidate such a dilemma. We found that Δ*M062R* infection failed to fine tune the translation machinery as observed during wildtype MYXV infection (**Figure 5A**). The obvious trend of sustained phosphorylation of 4EBP during wildtype MYXV infection is consistent with the behavior of wildtype VAC as previously reported.^30^ The effect by wildtype MYXV infection includes sustained activation of mTORC1 and p70S6 which signaling axis likely relieves 4EBP by hyper-phosphorylation; hypo-phosphorylated 4EBP-1 is an inhibition of the translation initiation complex.^31,32^ As controls, activities of mTORC1, p70S6, and 4EBP1 in the mock infected cells were also at a less degree than those during wildytype MYXV infection. During wildtype MYXV infection, sustained phosphorylation of AKT Threonine 473 is a hallmark for a permissive infection, which also is directly associated with MYXV oncolytic selectivity.^33,34,35,36^ However, Δ*M062R* infection did not activate AKT-Thr473 phosphorylation to the same degree as wildtype MYXV (**Figure 5B**).

**Figure 5.**
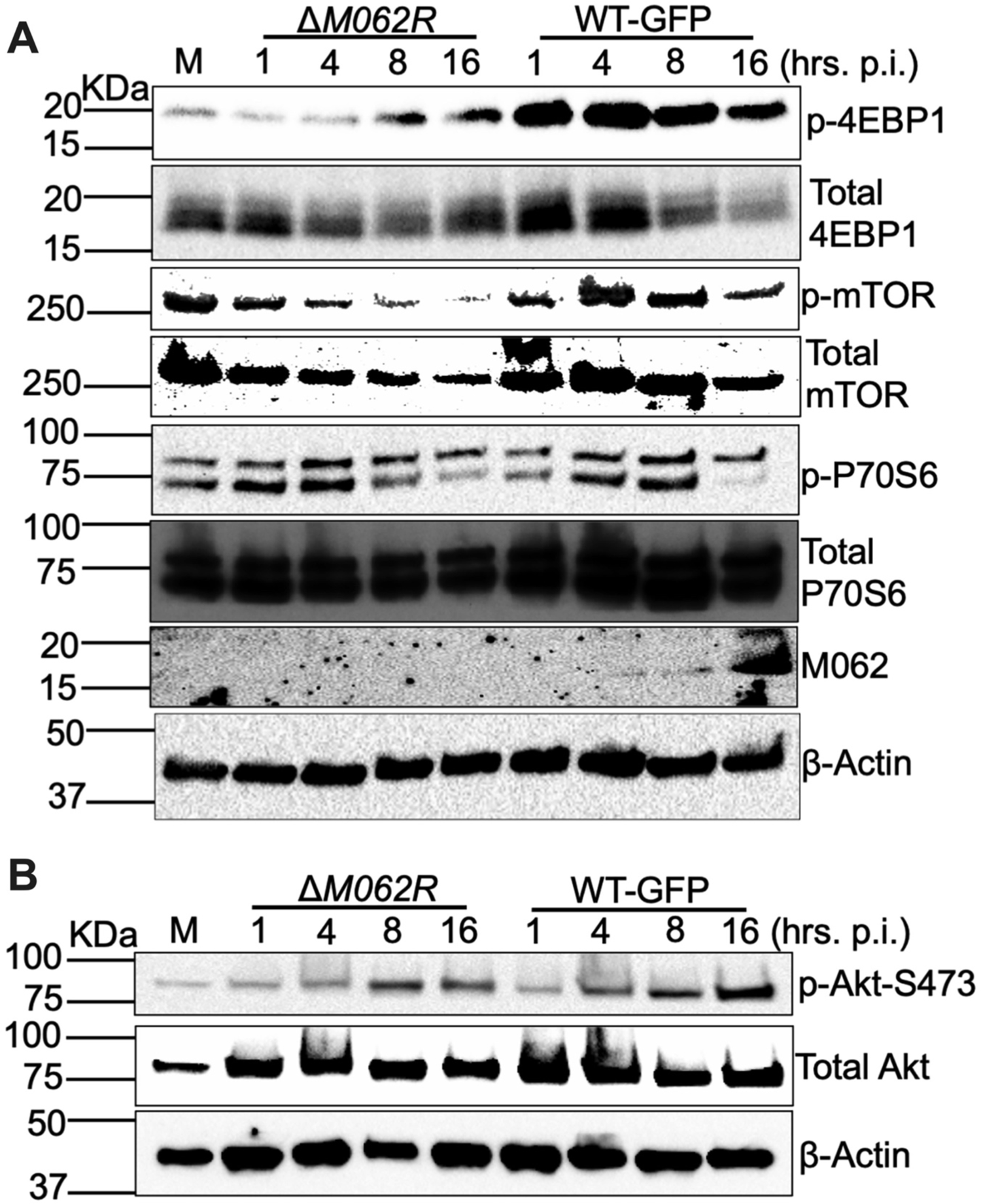
Infection by Δ*M062R* led to a distinct state of translation regulation in macrophages. In THP1 differentiated macrophages, different from wildtype MYXV Δ*M062R* infection did not lead to sustained activation of mTOR/P70S6/phosphor-4EBP1 (A) nor sustained phosphorylation of Akt-S473 (B). The time course experiments were performed and at given time points, equal amount of total protein lysates were assessed for signaling events.

We next examined the expression of ISGs in these macrophages on whether Δ*M062R* infection was associated with reduced host protein synthesis. We found on the contrary that many antiviral proteins are upregulated in expression compared to wildtype MYXV infection (**Figure 6A**). Many of these upregulated proteins are interferon-stimulated genes (ISGs) and their protein levels accumulated to be higher than those in the mock infection, e.g., MDA5, RIG-I, cGAS, PKR, and STING, etc. Similar phenomenon can be seen in primary human CD4+ monocytes/macrophages (**Figure 6C**) and primary human fibroblast (**Supplemental Fig 1**). Among proteins that we examined, some remain at the comparable expression levels with those in mock infection or wildtype MYXV infected cells (**Figure 6B**) suggesting that Δ*M062R* effect may apply to a subset of host protein. Overall, we did not observe specific downregulation of host protein expression during Δ*M062R* infection. Thus, although Δ*M062R* infection fails to cause host shut-off likely due to a lack of viral protein synthesis at the late stage of the infection, host protein synthesis in general, on the other hand, is unhindered. More importantly, in macrophages, Δ*M062R* treatment leads to upregulation of host proteins that play important roles in innate immunity and antiviral responses.

**Figure 6.**
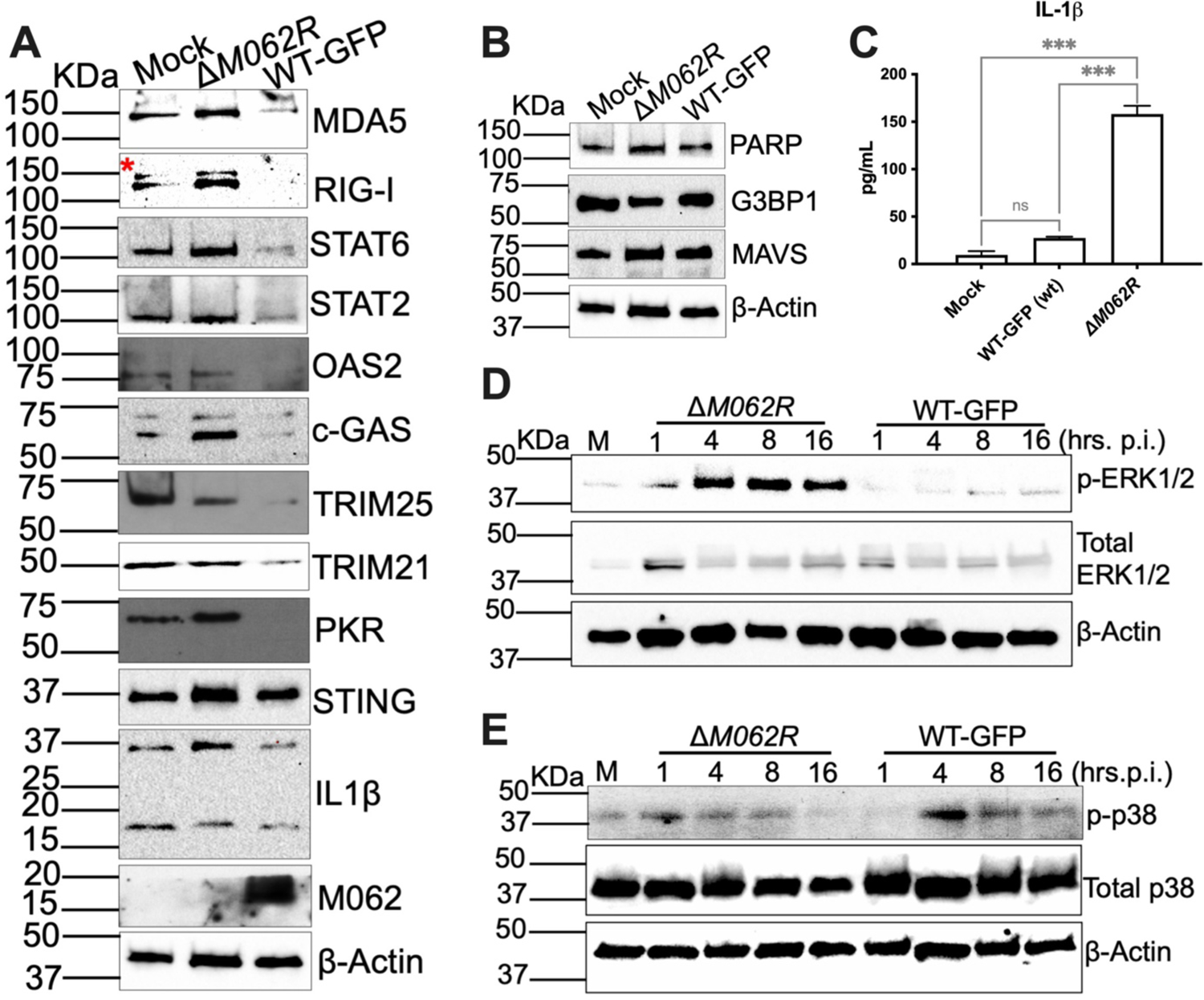
Treatment of Δ*M062R* in macrophages led to differential regulation of ERK1/2 and p38 signaling and upregulation of a subset of antiviral molecules. A. Differentiated THP1 cells were mock treated, infected with Δ*M062R*, or wildtype MYXV. At 24 hrs. post-treatment, numbers of host protein levels were compared among three groups. A subset of antiviral proteins were specifically upregulated compared to wildtype MYXV infection. B. In the same assay as in “A”, a cluster of host protein levels remained indifferent among three groups, mock treatment, Δ*M062R*, or wildtype MYXV infection. C. In primary healthy human CD14^+^ monocytes/macropahges, Δ*M062R* treatment also led to upregulation of IL1β detected by multi-plex immunoassay, which was consistent with what was seen in differentiated THP1 cells in “A”. D. Infection by Δ*M062R* led to activation of ERK1/2 signaling. E. While wildtype MYXV infection of THP1 macrophages promoted p38 activation, infection by Δ*M062R* attenuated such effect in p38 signaling.

### Treatment with Δ*M062R* induced a unique state in macropahges, which may promote heightened innate immune responses after exposure to new stimuli, such as dsRNA danger signals

Previously we found that the Δ*M062R* induced proinflammatory responses in the primary human macrophages.^20^ Treatment with this mutant virus also stimulated anti-tumor effect effectively compared to the oncolysis by the replication-competent wildtype MYXV.^37^ We found here that Δ*M062R* infection differentially impacted ERK1/2 and p38 activities (**Figure 6D** and **6E**). In contrast to wildtype MYXV infected cells, Δ*M062R* leads to activation of ERK1/2 signaling (phosphorylation) (**Figure 6D**) without activating p38 signaling (**Figure 6E**). This may correlate to the immunological consequences caused by these viral infections. For example, wildtype VACV promotes p38 activation in macrophages,^38^ which is consistent with what we observed during wildtype MYXV infection. Activating ERK1/2 signaling stimulates cell growth by activating numerous cellular transcription factors.^39,40^ This may explain our observation in RNAseq analyses that Δ*M062R* infection led to upregulation of cellular gene expression promoting cell survival and growth (**Supplemental Fig 2**).

Previously we reported that Δ*M062R* pretreatment followed by dsDNA exposure could lead to heightened IFN-I, while wildtype MYXV potently inhibited the effect.^20^ Because the expression of host innate immune modulators, such as RIG-I and cGAS, was intact or even increased after Δ*M062R* infection, we hypothesize that IFN-I induced by RNA sensors would be intact after Δ*M062R* infection, while wildtype MYXV infection could suppress host RNA sensing. We first tested serotype 3 reovirus (Reo TD3) as a stimulus, which is a known activator of both MDA-5 and RIG-I.^41^ We found that wildtype MYXV pretreatment effectively suppressed Reo TD3 stimulated IFN-I induction, while Δ*M062R* pretreatment promoted elevated IFN-I induction (**Figure 7A**). We next tested 5’ppp-dsRNA, a synthetic RIG-I agonist, for its ability to induce IFN-I; as a control, we used control 5’ppp-dsRNA without 5’-triphosphate moiety for the same test. We found that while wildtype MYXV again effectively suppressed the IFN-I induction, Δ*M062R* pretreatment potentiated the response to 5’ppp-dsRNA for having more effective IFN-I production (**Figure 7B**). On the other hand, control dsRNA did not induce IFN-I by itself, while significant elevation of IRF-driven gene expression in Δ*M062R* group was observed that is due to activation of DNA sensing pathway by Δ*M062R* infection (**Figure 7C**). This outcome is consistent with what we reported previously.^20^

**Figure 7.**
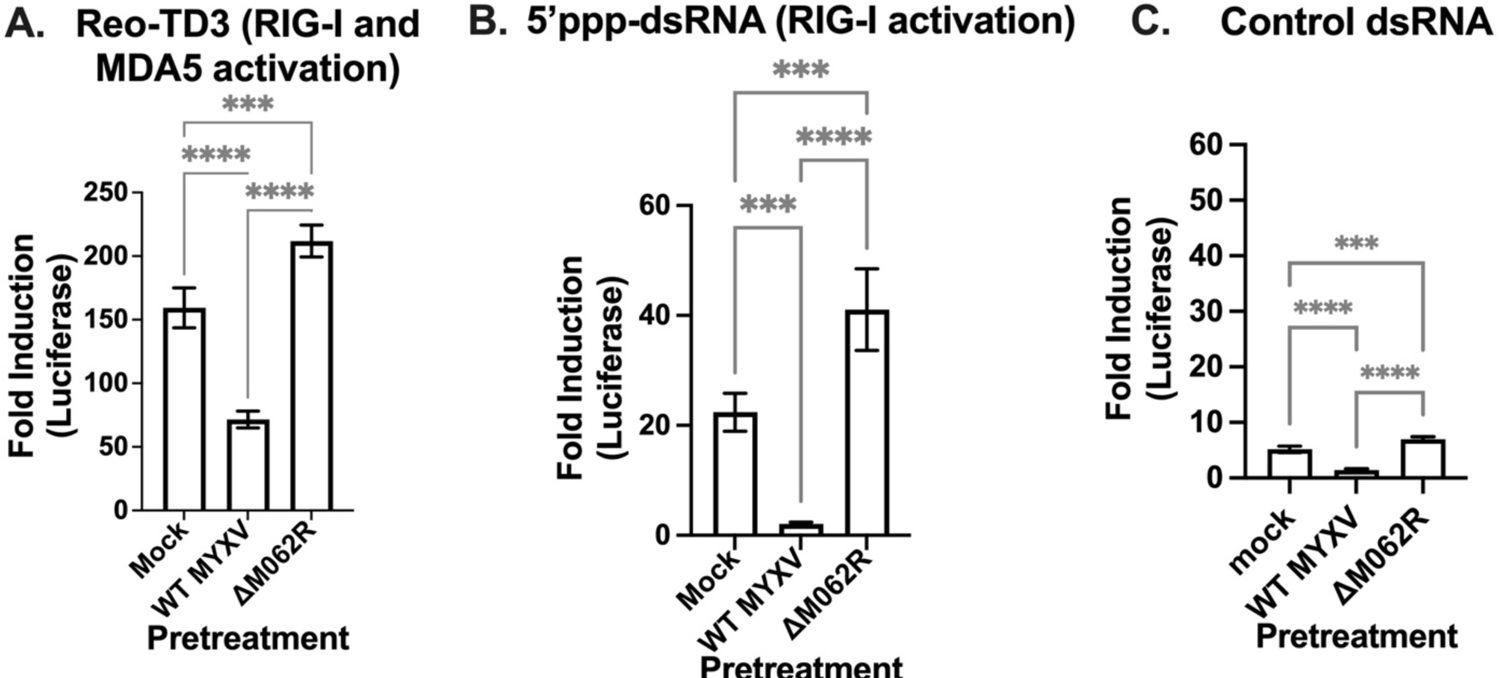
Pretreatment of Δ*M062R* in macrophages potentiated the IFN-I responses when they were exposed to new innate immune stimuli. **A. While** pretreatment of macrophages with wildtype MYXV significantly inhibited Reo TD3 induced IFN-I induction indicated by IRF-dependent luciferase expression, Δ*M062R* pretreatment led to heightened responses. Reo TD3 is a known stimulus to both RIG-I and MDA5 for IFN-I induction. B. Pretreatment of macrophages with Δ*M062R* led to heightened IFN-I induction when cells were exposed to a new rounds of RIG-I specific stimuli (5’ppp-dsRNA). On the contrary, wildtype MYXV potently suppressed RIG-I activation by 5’ppp-dsRNA. C. As control, the 5’ppp-dsRNA without the 5’-triphosphate moiety was used for a similar test as in “B”. The control dsRNA by itself did not induce IFN-I, while Δ*M062R* induced IFN-I significantly due to activation of DNA sensing pathway that we reported previously. Statistical analyses of ordinary one-way ANOVA followed by Tukey’s multiple comparison test were performed. ***p<0.0005, ****p<0.0001.

## Discussion

Poxviruses evade host defense with multi-faceted strategies, and thus are excellent toolbox for the understanding of cell biology and immunity. Myxoma virus (MYXV) M062 protein belongs to the poxvirus host range *C7L* superfamily.^9^ The host target of MYXV M062 is SAMD9 and antagonizing SAMD9 is so far the only functional relevance observed.^10,11,26,20^ The direct interactions between MYXV M062 and host SAMD9 at the SAMD9 AlbA_2-like domain^26^ suggests its potential importance to inhibit SAMD9 from binding to double stranded nucleic acids (dsNA) and performing nuclease activities.^13^ This hypothesis was based on recent finding that the AlbA_2-like domain had binding affinity to dsNA^21^ and nuclease activities against tRNA^phe^.^13^ However, the role of SAMD9 in cell biology or immune responses remains to be understood, which will be the key to understand the complex human diseases caused by deleterious mutations in SAMD9.^42,17,19^ Because so far MYXV M062 is the only poxvirus protein shown to directly interact with SAMD9,^26^ it is a key tool to reveal the possible regulatory mechanism of SAMD9. The loss of viral *M062R* gene during infection could unavoidably activate SAMD9 function;^20^ it will be true especially if M062 targeting domain of SAMD9 is important for the on/off switch of SAMD9 antiviral property. In this study, we thus examined the defect in infection caused by the mutant MYXV with targeted deletion of *M062R* gene and examined its immunological effects and utility. More importantly, with the pending application of using this mutant virus as an immunotherapeutic platform, the immunological impact induced by treatment of Δ*M062R* needs to be understood.

Infection by Δ*M062R* in most cells tested is abortive. We found it intriguing that despite its inability of late viral protein synthesis and producing progeny virions, viral RNA synthesis including intermediate and late RNAs remains competent. The main defect observed during Δ*M062R* infection is the severely attenuated viral DNA replication and late protein synthesis. Surprisingly, intermediate protein synthesis during Δ*M062R* infection remains competent, albeit moderate decline in levels compared to the wildtype infection. This is an interesting phenomenon, which suggests, at least in the cell types we investigated, that viral DNA genome amplification may have triggered SAMD9’s ability to recognize danger signal and to activate its antiviral function. In that, the activated SAMD9 may specifically inhibit translation of viral mRNA but release the inhibition or promote the synthesis of certain host proteins for the elimination of the threat. What is the danger signal to SAMD9 and how it differentially acts on danger or self mRNA remain to be understood.

Phylogenetic analyses revealed M062’s divergency to highly conserved orthopoxvirus *C7L* homologs, designated Clade I members.^9^ Although VACV C7 protein appeared binding to SAMD9, it may have evolved to target other pathways connecting to SAMD9 pathway (unpublished data and ^43^). This is not surprising, as mammalian poxviruses employ different strategies to evade host surveillance and defense mechanisms, such as lessons learned from poxvirus inhibition of NF-κB signaling.^44^ In this study, we revealed the unique translation state induced by Δ*M062R*, which is different from what was observed during infection by VACV-Δ*C7L*Δ*K1L*, the mutant VACV without both inhibitors of SAMD9.^29^ Although Δ*M062R* viral infection is abortive, we did not observe abnormality in host protein synthesis and accumulation.

Furthermore, we detected upregulated protein levels for a subgroup of factors comprising antiviral and pro-inflammatory proteins after Δ*M062R* infection in both macrophages and primary cells. This suggests a previously uncharacterized mechanism in distinguishing viral and host self to trigger host defense through calculatingly promoting *de novo* protein synthesis of antiviral molecules. More importantly, Δ*M062R* pretreatment potentiated macrophages for heightened proinflammatory responses against new danger stimuli including cytoplasmic dsDNA and dsRNAs. The potential link between transcriptomic rewiring and translation reprograming converges on the activity of effect induced by Δ*M062R* infection, which urges further investigation on the mechanism. Because Δ*M062R* showed promising therapeutic benefit in subverting immunosuppressive myeloid cells (unpublished data), understanding the mechanism of Δ*M062R* induced cellular state is critical to development of novel therapy against immunological conditions including cancer and improvement of vaccine efficacy through reprograming innate and intrinsic immune responses.

## Material and Method

### Cell culture and virus stock

Mammalian cells used in this study include BSC-40 (ATCC CRL-2761) (in Dulbecco minimal essential medium, DMEM, Lonza), HeLa (ATCC CCL-2 cultured in DMEM, Lonza), HEK293T (ATCC CRL-3216, DMEM), THP1 (ATCC TIB-202), and THP1-dual Lucia (Invivogen) or THP1-Lucia (kindly provided by F. Zhu Florida State University)^45^ (in RPMI1640, Lonza). Primary normal human CD14^+^ monocytes (Lonza, Walkersville, MD) (RPMI1640) and primary normal dermal fibroblasts NHDF-neo (Lonza, Walkersville, MD) (5% FBS-DMEM) were also used in this study. The complete growth medium (e.g., DMEM Lonza/BioWhittaker Catalog no 12-604Q, or RMPI1640) was supplemented with 10% FBS (Atlanta Biologicals, Minneapolis, MN) unless specified otherwise, 2 mM glutamine (Corning Cellgro, Millipore Sigma, St. Louis, MO), and 100 μg per ml of Pen/Strep (Corning Cellgro, Millipore Sigma, St. Louis, MO); for RPMI1640 complete culture medium, in addition to FBS, glutamine, and Pen/Strep, 2-mercaptoethanol (MP biomedicals, Solon, OH) was supplemented to a final concentration of 0.05 mM.

The viruses used in the study are from myxoma virus (MYXV) Lausanne strain background (GenBank Accession AF170726.2).^10^ The *M062R*-null MYXV (Δ*M062R*) targeted deletion mutant and wildtype virus (WT-GFP) were described previously.^10^ Myxoma virus stocks were amplified on BSC-40 cells and purified with sucrose step gradient through ultracentrifugation as previously described.^46^ Reovirus TD3 was provided by K. W. Boehme (University of Arkansas for Medical Sciences) and described previously.^41^

### Realtime PCR to examine viral DNA replication

Cells were infected with either wildtype MYXV (WT-GFP) or Δ*M062R* at an MOI of 5. At the given time points (4, 8, 12, 24 hrs. p.i.) DNA extraction was performed using DNeasy Blood & Tissue Kit (Qiagen, Cat. No. 69506) followed by realtime PCR (Luna Universal qPCR Master Mix, Cat. No. M3003, NEB Inc) following manufacturer standard protocol. Realtime PCR primers recognizing *M071L* were used for quantification of viral DNA levels as previously described.^10^

### Luciferase assay to examine viral intermediate gene expression

HeLa cells were transfected with the plasmid containing the luciferase expression cassette driven by one of following VACV intermediate gene promoters, pA2L,^24^ pI1L,^24^ and VACV intermediate consensus promoter sequence (pINT)^25^ (M. Wiebe, University of Nebraska-Lincoln) using ViaFect (Cat# E4982, Promega, Madison, WI) at 0.8 μg/mL with the transfection agent to DNA ratio of 3 μL to 1 μg as previously described.^20^ At 6 hours post-transfection, cells were in the presence of 200 μM Ara-C (Sigma-Aldrich, C6645) infected with either wildtype or Δ*M062R* MYXV at a MOI of 3 for 16 hours before harvesting with Passive Lysis Buffer (Cat# 1910, Promega) and measuring luciferase activities using Luciferase Assay System (Cat# E1500, Promega) following manufacturer instructions.

### Western blot and Multi-plex Cytokine Array

For THP-1 cells, 10^6 cells were differentiated with PMA (FisherScientific, Cat# BP685-1) at 50 ng/ml for up to 48 hours (hrs) before mock treated, infection with wildtype, or Δ*M062R* at an MOI of 5. Cells were harvested at given time points of 1, 4, 8, 16 hrs post-infection (p.i.) with the lysis buffer [1% NP-40, 50mM Tris-Cl pH 7.4, 150mM NaCl containing the protease inhibitor (cOmplete ULTRA EDTA-free Protease inhibitor, Cat# 6538282001, MilliporeSigma), 2 mM sodium orthovandate (Santa Cruz Biotechnology, Inc, Cat#13721-39-6), 1 mM PMSF (Cat# 52332, MilliporeSigma), and 25mM β-Glycerophosphate (SigmaAldrich, Cat# G9422)]. Cell lysates were incubated on ice for 30 minutes before centrifugation at 125,000 g for 10 min and 50 μg of total protein for each sample was separated on SDS-PAGE [Mini-PROTEAN TGX Gels (Biorad, Cat# 456-1095)]. Proteins were transferred to 0.2um Nitrocellulose membrane (Cat# 1620112) for antibody probing. Antibodies used in this study include mTOR (Cell Signaling Technology or CST, Cat# 2983P), Phospho-mTOR Ser2448 (CST, 5536S), p70S6 Kinase (CST, 9202S), Phospho-p70S6 Kinase T421/S424 (CST, 9204S), 4EBP1 (CST, 9644S), Phospho-4EBP1 (CST, 9456S), AKT (CST, 4961S), Phospho-AKT S473 (CST, 4060S), Phopsho-P44/42-MAPK T202/204 (CST, 4370S), MAPK P44/42 (CST, 4695S), Phospho-p38 T180/Y182 (CST, 9216S), p38 (CST, 9212S), β-Actin (Sigma-Aldrich, A1978), rabbit anti-M062, M038, and SERP-1 antibodies were generated by BIOMATIK (Ontario, Canada). For detection of IL1β, the Human Inflammation Panel (Invitrogen™ Inflammation 20-Plex Human ProcartaPlex™ Panel) (Catalog # EXP20012185901, eBioscience/Invitrogen) was used following manufacturer standard protocol using BioRad Bio-Plex 200 system (Bio-Rad, Hercules, CA).

### Pulse-labeling to detect newly synthesized proteins

Differentiated THP1 cells were infected with wildtype or Δ*M062R* MYXV at an MOI of 5 to 10 for 12 hrs before culturing in medium free of methionine and cystine (Gibco, Cat# 21013024) for 1 hr. Pulse-labeling was performed with 25 µM AHA (L-Azidohomoalanine) (Invitrogen, Cat# 10102) for 2 hrs before cell lysate harvesting in 120 µl of RIPA lysis buffer (1% NP-40, Tris pH 7.4, NaCl 150mM, 0.5% Sodium deoxycholate, 0.1% Sodium dodecyl sulfate, and theprotease inhibitor). Cell lysates were treated with 250U of Benzonase (Novagen, Cat# 70664-3) and incubate for 30 minutes before centrifugation at 12,500 g 4°C for 10 min, 200 µg total proteins were subjected to Click-iT chemistry reaction using 100 µL of Click-iT Reaction Buffer (Invitrogen, Cat# 10276) and 40 µM of Biotin Alkyne (Invitrogen, Cat# B10185) following manufacturer’s instructions. After precipitation (methonal and chloroform) 20 µg of resolubilized protein (1% SDS in 50 mM Tris-HCl pH8.0) from each sample was analysed in Western blot probed with Streptavidin-HRP (Thermo scientific, Cat# 21130).

### Next Generation RNA sequencing

THP-1 cells (10^6^ cells per 3.5 cm dish) were differentiated in PMA at 50 ng/mL for up to 48 hours before mock treated, transfection with interferon stimulated DNA (ISD)^47^ at 2 μg per dish using ViaFect based on manufacturer protocol (Cat. No. E4982, Promega, Madison, WI), or infected at an moi of 5 for either wildtype MYXV (WT-GFP) or Δ*M062R*. At 1-, 4-, and 8-hours post-transfection of ISD and 1-, 4-, 8-, and 12-hours post-infection cells were harvested and spun down for RNA extraction. RNA extraction was performed as previously described.^20^ Briefly, after Quick DNA/RNA MiniPrep Plus kit (Cat. No. D7003; Zymo, Irvine, CA, USA), quality control of purified RNA was performed using the Qubit RNA BR Assay kit (Cat. No. Q10211; Invitrogen, Waltham, MA, USA) and the Standard Sensitivity RNA Analysis kit on a Fragment Analyzer capillary electrophoresis system (Cat. No. DNF-471-0500; Agilent, Santa Clara, CA, USA).

TruSeq Stranded Total RNA library prep kit with unique-dual indexing (Cat. No. 20020598 & 2002371; Illumina, San Diego, CA, USA) was used for this study with 250 ng of total RNA. Quality control of the libraries was conducted using the Qubit 1X dsDNA HS Assay kit (Cat. No. Q33231; Invitrogen, Waltham, MA, USA) for mass, the High Sensitivity NGS Fragment Analysis kit on a Fragment Analyzer capillary electrophoresis system (Cat. No. DNF-474-0500; Agilent, Santa Clara, CA, USA) for fragment size, and the Universal Library Quantification kit (Cat. No. 07960140001; KAPA, Wilmington, MA, USA) for functional validation. Finally, validated libraries were adjusted to 3 nM followed by pooling, denaturing, and clustering. HiSeq 3000 (Illumina, San Diego, CA, USA) was used for paired end (2X75) sequencing to an average of 40 million reads per sample.

### Dual-RNAseq bioinformatic data processing and GO:BP analyses

A dual-RNAseq bioinformatic data processing pipeline was utilized for the study as previously described.^20^ Exploratory data analysis (EDA) and differential expression analysis was performed using R and DESeQ2 v1.32.0, where p-value and adjusted p-value thresholds were set to 0.05 and 0.1, respectively. Gene expression comparisons within figures were made by normalizing raw count data using the median of ratios method in DESeQ2.^23,22^ The Reactome pathway enrichment plot were generated using the R library ReactomePA v1.44.0 and the *enrichPathway* function.^48^ Only upregulated significant differentially expressed genes were passed to the aforementioned function and the top 10 pathways are displayed. GO term (biological processes) heatmaps and enrichment analysis was performed using the R library gProfileR v0.7.0.^49^ All significant differentially expressed genes were passed for analysis, and “strong” hierarchical clustering was chosen. For heatmap generation, the mean DESeq2 normalized expression values for those genes overlapping a given GO:BP terms was used.

### Luciferase based assay to assess type 1 interferon activation

THP1 cells with IRF core promoter driven luciferase expression and in some cases THP1-dual cells (IRF-luciferase and NF-κB-SEAP, Invivogen, Cat. No. thpd-nfis) were differentiated into macrophages as previously described.^20^ After pretreatment of mock infection, infection with WT MYXV or Δ*M062R* at an MOI of 5 for 9 hours before exposure to reovirus TD3^41^ at an MOI of 50. Other experiments were performed by transfection of 5’ppp-dsRNA (RIG-I activator) (Invivogen, Cat. No. Tlrl-3prna) with Lipofectamin RNAiMAX (Invitrogen, Cat. No.13788-075) after the pretreatment. At 12 hours post the secondary infection or exposure to stimuli such as 5’ppp-dsRNA, supernatant was harvested for luciferase assay (QUANTI-Luc, Invivogen, Cat. No. rep-qlc1) following manufacturer’s standard protocol. As a control for 5’ppp-dsRNA, the non-stimulatory dsRNA to RIG-I that does not contain the 5’-triphosphate moiety (Invivogen, Cat. No. Tlrl-3prnac) was also included in the experiment.

### Statistical analyses

GraphaPrism 10.1.1 was used for statistical analyses. One-Way ANOVA followed by individual group comparisons (i.e, multiple comparison followed by Ordinary One – Way ANOVA) or Two – way ANOVA, and Student t test or multiple unpaired t test was performed according to experiment set up and statistical significance is defined as *p<0.05 (**p<0.01, ***p<0.001).

### Data submission

Sequence data were deposited at the NCBI Gene Expression Omnibus (GEO) and will be released upon publication of the manuscript.

## Acknowledgements

The study initiated in 2016 was supported by NIH K22AI099184 and R01AI139106 to JL* (corresponding author), a fund from the River Valley Ovarian Cancer Coalition, a start-up by UAMS Department of Microbiology and Immunology, a UAMS Barton pilot award, and a UAMS ABI-VCRI Collaborative award to JL*. This work is also supported in part by the Center for Microbial Pathogenesis and Host Inflammatory Responses grant P20GM103625 through the NIH National Institute of General Medical Sciences (NIGMS) Centers of Biomedical Research Excellence (COBRE) and by the Winthrop P. Rockefeller Cancer Institute at UAMS to JL*. We thank Sarah Blair, Bernice Nounamo, Shana Chancellor, and Richard Connor for technical support.

---

**Supplemental Fig 1.**
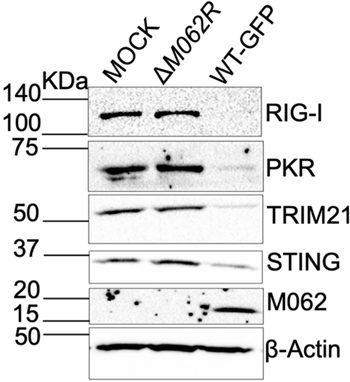
Infection by Δ*M062R* in healthy primary human dermal fibroblasts showed intact host protein synthesis. Target protein levels from mock infected, infected with Δ*M062R*, or wildtype MYXV were compared in primary NHDFneo. At 24 hrs. post-treatment, Western blot was performed. Infection by wildtype MYXV led to depletion of host proteins tested, while Δ*M062R* did not hinder these protein levels.

**Supplemental Fig 2.**
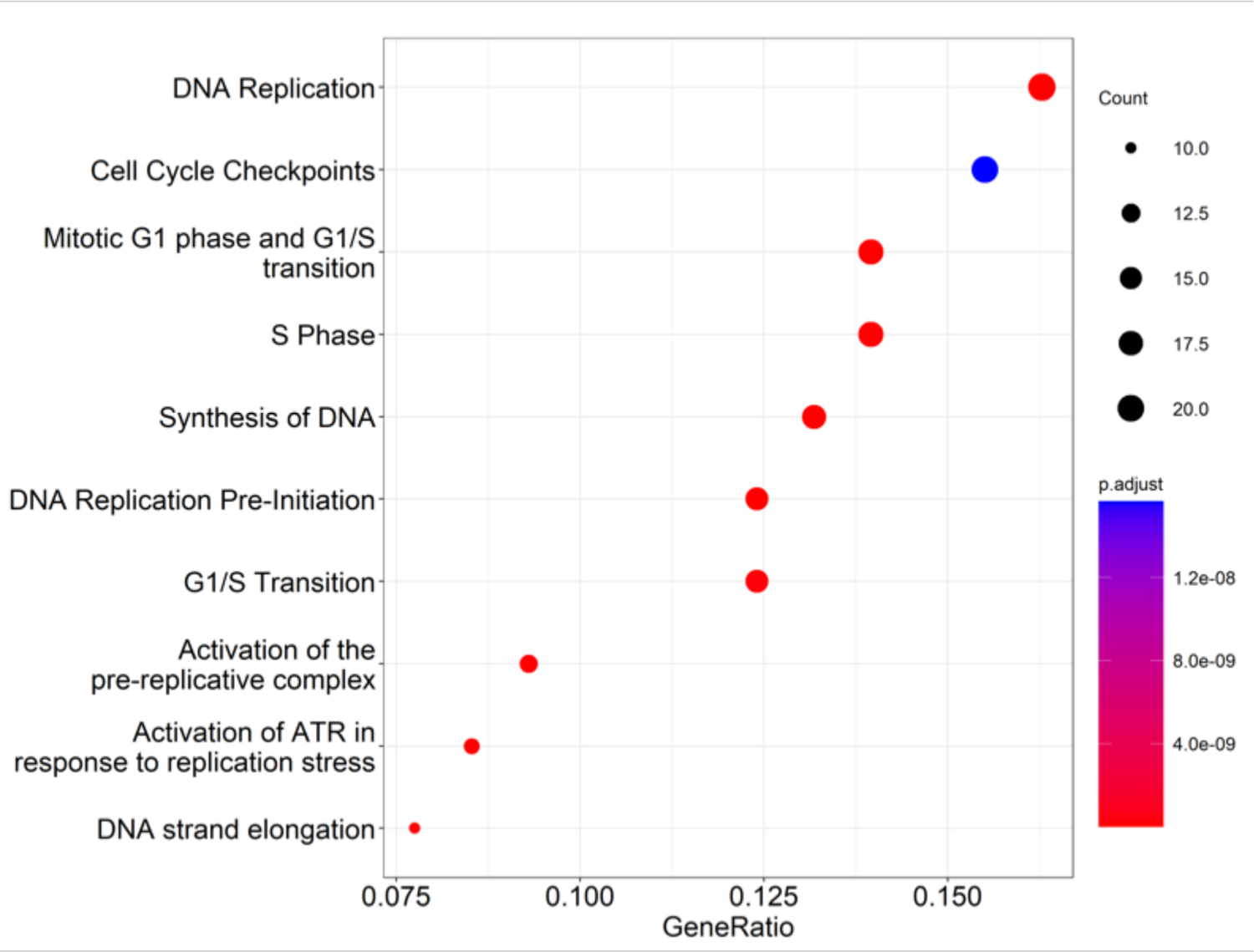
Treatment by Δ*M062R* in human macrophages led to unique state in these cells promoting survival and growth compared to wildtype MYXV. At 12 hrs. p.i., compared to wildtype MYXV treatment, Δ*M062R* induced gene expression determined by GO biological processes enrichment analysis of differentially expressed genes from RNAseq data analyses. The left y-axis and upper x-axis display the dendrogram resulting from the hierarchical clustering of the Euclidean distance between samples and GO:BP terms. A heatmap displays the z-score standardized expression of the genes within a given term, blue for lower normalized expression and red for higher. GO:BP terms on the right y-axis are encoded red and green, for induced and repressed terms, respectively. Sample names are displayed along the lower-x-axis.

